# Murine Neutrophil Chemotaxis Following Burn Injury with Poloxamer 188 Treatment in a Microfluidic Platform

**DOI:** 10.64898/2026.01.07.698204

**Authors:** Hossein Razmi Bagtash, Nagham Alatrash, Udaya Sree Datla, Bhaskar Vundurthy, Shuai Shao, Roopa Koduri, Mecca Islam, Kevin Mutore, Emma Salari, Renqing Wu, Vanessa Nomellini, Caroline N. Jones

**Affiliations:** Department of Bioengineering, University of Texas at Dallas, Richardson, TX, USA; Department of Biomedical Engineering, UT Southwestern Medical Center, Dallas, TX, 75390, USA; Division of Burn, Trauma, Acute, and Critical Care Surgery, Department of Surgery, UT Southwestern Medical Center, Dallas, TX, USA; Department of Biomedical Engineering, Carnegie Mellon University, Pittsburgh, PA, USA; Robotics Institute, Carnegie Mellon University, Pittsburgh, PA, USA; Department of Biology, University of Texas at Dallas, Richardson, TX 75080, USA

**Keywords:** Poloxamer 188 (P188), neutrophil chemotaxis, microfluidic system, burn injury

## Abstract

This study investigates the effects of Poloxamer 188 (P188) on neutrophil chemotaxis following burn injury in male and female mice using a microfluidic system. Utilizing male and female CD1 mice, we evaluated neutrophil migration towards two chemoattractants, N-formyl-l-methionyl-l-leucyl-l-phenylalanine (fMLP) and Leukotriene B4 (LTB4), and FPR1 and BLT1 G protein-coupled receptors after administering P188. Our findings revealed that P188 significantly increased the migration toward LTB4 in both sexes. Additionally, our findings highlight the upregulation of BLT1 and FPR1 markers due to burn injury in both female and male mice in the Burn vs. Sham groups. These results demonstrate the potential of P188 in modulating neutrophil behavior post-burn injury in therapeutic strategies for inflammation management. This microfluidic platform offers a precise and controlled microenvironment for studying neutrophil chemotaxis post-burn injury with and without P188 treatment.

## 1. Introduction

In severe burn injury, the local inflammatory reaction leads to secondary organ damage by triggering excessive cytokines release^1–3^. Poloxamer 188 (P188), a synthetic and amphiphilic FDA-approved copolymer^4^, integrates into the damaged cell membranes thereby reducing the cell permeability and preventing further cell damage^5–7^. Recent studies show the effect of P188 on regulating inflammation and alleviating the detrimental effects of systemic inflammation and secondary organ injury in burn and other inflammatory conditions, which significantly impedes recovery^8–11^. In addition to its membrane-stabilizing properties, P188 has been shown to modulate neutrophil activation and function in a dose-dependent manner. Harting et al. reported that P188 influences neutrophil oxidative burst and CD11b expression, where low concentrations enhance oxidative activity, while higher concentrations can attenuate both oxidative burst and surface CD11b expression in activated neutrophils^9^. This finding indicates that P188 not only protects cellular integrity but also regulates immune cell activation, providing a dual mechanism that may mitigate excessive inflammatory responses following severe burn injury. To further evaluate these immunomodulatory effects in vivo, animal models offer a controlled platform to study systemic inflammation and immune cell behavior in response to injury and therapeutic interventions. Mice models are widely used in a variety of research areas, particularly inflammation and burn injuries^12–16^. In response to burn injury, damage associate molecular patterns (DAMPs) are secreted which lead to inflammation and consequently result in neutrophil activation and migration to the site of damage to clear the pathogen and secret pro-inflammatory mediators^12,17^. With this, however, uncontrolled and dysfunctional migration of neutrophils can worsen inflammation and impede the healing process^18–21^.

There is growing evidence correlating sex differences with mortality and clinical outcomes in burn injuries^22^. Among studies on burn injuries, the majority of them show that women are at higher risk of mortality when sustaining similar injuries ^22,23^, but the reason is still not fully understood. It is hypothesized that sex hormones like estrogen can result in immune system dysregulation and worsen the outcomes in women^24–28^. We controlled for the main factors, including age, burn size, and lack of inhalation damage to study the impact of sex difference and P188 drug in burn and immune system function. To better understand the potential sex hormone impact on neutrophil migration, better tools are needed^29^. Traditional assays often fail to capture subtle, dynamic changes in neutrophil behavior; therefore, integrating advanced technologies becomes essential for uncovering these mechanisms.

Microfluidics are novel technologies that enable precise measurement of neutrophil migratory behavior and can offer new insight into neutrophil chemotaxis and their role in the inflammatory response after burn injury^30–32^. Further studies on the precise measurement of neutrophil migration may lead to the development of new interventions to regulate the overwhelming inflammatory response in burn patients^33–36^. Chemotaxis, which is guided cell migration toward chemoattractant signals, plays an indisputable role in recruitment of neutrophils to the sites of inflammation^37–41^. Neutrophils prioritize among signals they receive, and there is a hierarchical order in response to signaling cascades neutrophils receive^17,42–44^. Multiple chemoattractants stimulate neutrophil chemotaxis in which N-formylated peptides like N-formyl-l-methionyl-l-leucyl-l-phenylalanine (fMLP) and lipid chemoattractants like leukotriene B4 (LTB4) have been studied in this field. In this research, we reported the fMLP/LTB4 percent neutrophil migration ratio for murine neutrophil responsiveness to both signals. We believe this ratio provides a straightforward metric to better interpret neutrophil responsiveness to these stimuli. Additionally, the ratio simplifies the data into a single value that reflects the sensitivity of neutrophils to both bacterial chemoattractant (fMLP) and pro-inflammatory (LTB4) signals^45^.

In this study we evaluated the effects of Poloxamer 188 (P188) on neutrophil chemotaxis following burn injury using both in vivo and ex vivo experiments in both male and female mice. We examined the impact of P188 on four distinct groups of male and female CD1 mice (sham, sham + P188, burn, and burn + P188) to study their migration behavior towards the chemoattractants N-formyl-l-methionyl-l-leucyl-l-phenylalanine (fMLP) and Leukotriene B4 (LTB4). In addition to evaluating neutrophil migration, we assessed the expression of CD11b, FPR1, and BLT1 markers to measure neutrophil activation, and we quantified reactive oxygen species (ROS) production across the four experimental groups. Our findings revealed that P188 significantly enhanced migration toward LTB4 in both sexes. Furthermore, our results highlight the upregulation of BLT1 and FPR1 markers due to burn injury, with significant increases observed in both male and female burn vs. sham groups.

## 2. Materials and Methods

### 2.1 Severe burn injury mouse model

Male and female CD1 mice at 11-12 weeks old were anesthetized, hair was removed from their back, and mice either underwent a sham injury or metal brands were used to create a 30-40% total body surface area burn injury. For sham injury mice underwent the same procedures as burn, including anesthesia and hair removal, but sham injury mice didn’t receive the actual burn, being as a control group to study any effects unrelated to the burn injury itself. Both sham and burn-injured mice were given 4 mL/kg body weight/total body surface area % of Lactated Ringer’s solution for resuscitation and analgesia was given intraperitoneally with buprenorphine. One set of mice received vehicle, and another set received 15 µg/mouse Poloxamer-Alexa Flour 647 (P188-AF) intravenously (IV) through the tail vein one hour after injury. The mice were then sacrificed 24 hours after burn injury. Experimental groups were categorized as follows: 1) sham + vehicle, 2) sham + P188, 3) burn + vehicle, and 4) burn + P188. After sacrificing, blood samples were collected into blood collection tubes with EDTA (BD 367856, Fisher Scientific) to evaluate the effect of P188 on neutrophil chemotaxis in our microfluidic system.

### 2.2 Drug Conjugation and Administration

Poloxamer 188 (P188) was conjugated with Alexa Fluor 647 to visualize tissue distribution. Male and female CD1 mice were subjected to either a sham procedure or a 40% total body surface area (TBSA) using a metal brand. At 1 h post-injury, animals received a tail vein injection of either vehicle or Alexa Fluor–conjugated P188 (15 µg per mouse).

### 2.3 Tissue Collection and Processing

At 24 h post-injection, mice were euthanized, and major organs including skin, liver, kidney, lung, and spleen were harvested. Organs were then placed in 10% neutral buffered formalin (Fisher Scientific, Hampton, NH, USA ) for 48 hours. Samples were then placed in 10% sucrose overnight in 4°C. They were then transferred to 18% sucrose overnight in 4°C. Tissue was washed with 1x PBS (Gibco, Waltham, MA, USA) before being placed in cryomolds. Cryomolds were filled with OCT compound (Sakura Fintek, Torrance, CA, USA) to fully cover tissue. Lastly, samples were cryosectioned at 20 µm thickness and mounted on glass slides. Coverslips were mounted using DAPI mounting media (Abcam, Cambridge, UK) to visualize nuclei.

### 2.4 Imaging

Tissue sections were examined using a laser scanning confocal microscope (Fluoview FV3000, Olympus, Tokyo, Japan) and visualized using Fluoview software (FV315-SW, Olympus, Tokyo, Japan). Alexa Fluor fluorescence was detected at excitation/emission peak of 650-750 nm, and DAPI was detected at excitation/emission peak of 430-470 nm was imaged to identify nuclei.

All animal experiments were conducted in accordance with the Guidelines for the Care and Use of Laboratory Animals and were approved and supervised by the Institutional Animal Care and Use Committee (IACUC) at the UT Southwestern Medical Center.

### 2.5 Neutrophil isolation for chemotaxis assay

Neutrophils were isolated from the mice blood samples using the EasySep™ Mouse Neutrophil Enrichment Kit (STEMCELL Technologies, Cambridge, MA). In summary, the whole blood and lysis buffer (Ammonium Chloride Solution, STEMCELL Technologies) were mixed at a 1:9 ratio and incubated on ice for 15 minutes. Then the sample was centrifuged, and after removing the supernatant, the lysed cells were washed with EasySep buffer (STEMCELL Technologies). After the lysis step, the Manual EasySep™ Protocol was followed to isolate neutrophils. The isolated neutrophils were then stained with 20 µM of Hoechst stain (Thermo Fisher Scientific) for 10 minutes at 37 °C and 5% CO2. Finally, the stained neutrophils were resuspended at a concentration of 10^7^ cells/mL in IMDM+20% FBS to load into the cell loading chamber.

### 2.6 Microfluidic chemotaxis device design and fabrication

The microfluidic chemotaxis device used to study murine neutrophil migration consists of two chemoattractant chambers on opposite ends and a central cell loading channel. Each chamber has its own inlet and outlet (Fig. 1). Perpendicular migration channels connect the cell loading chamber to chemoattractant chambers, providing the possibility of directional migration. For master wafer fabrication, traditional photolithography techniques were used with a Mylar photomask (FineLine Imaging, Colorado Springs, CO, USA). Polydimethylsiloxane (PDMS)-based microfluidic devices were then made by pouring a 10:1 ratio of PDMS base and a curing agent by casting from the master wafer. To remove the bubbles, the master wafer was left inside the desiccator for 30 mins. After curing for 24 h in an oven at 65°C, the PDMS mold was peeled off the master wafer, and inlet and outlet ports were punched using 0.75 mm punchers (Ted Pella, Reading, CA, USA). Finally, the punched PDMS-based devices, and 24-well glass-bottom plate were treated using Harrick plasma cleaner and bonded together on a hotplate at 80°C for 10 mins.

**Figure 1.**
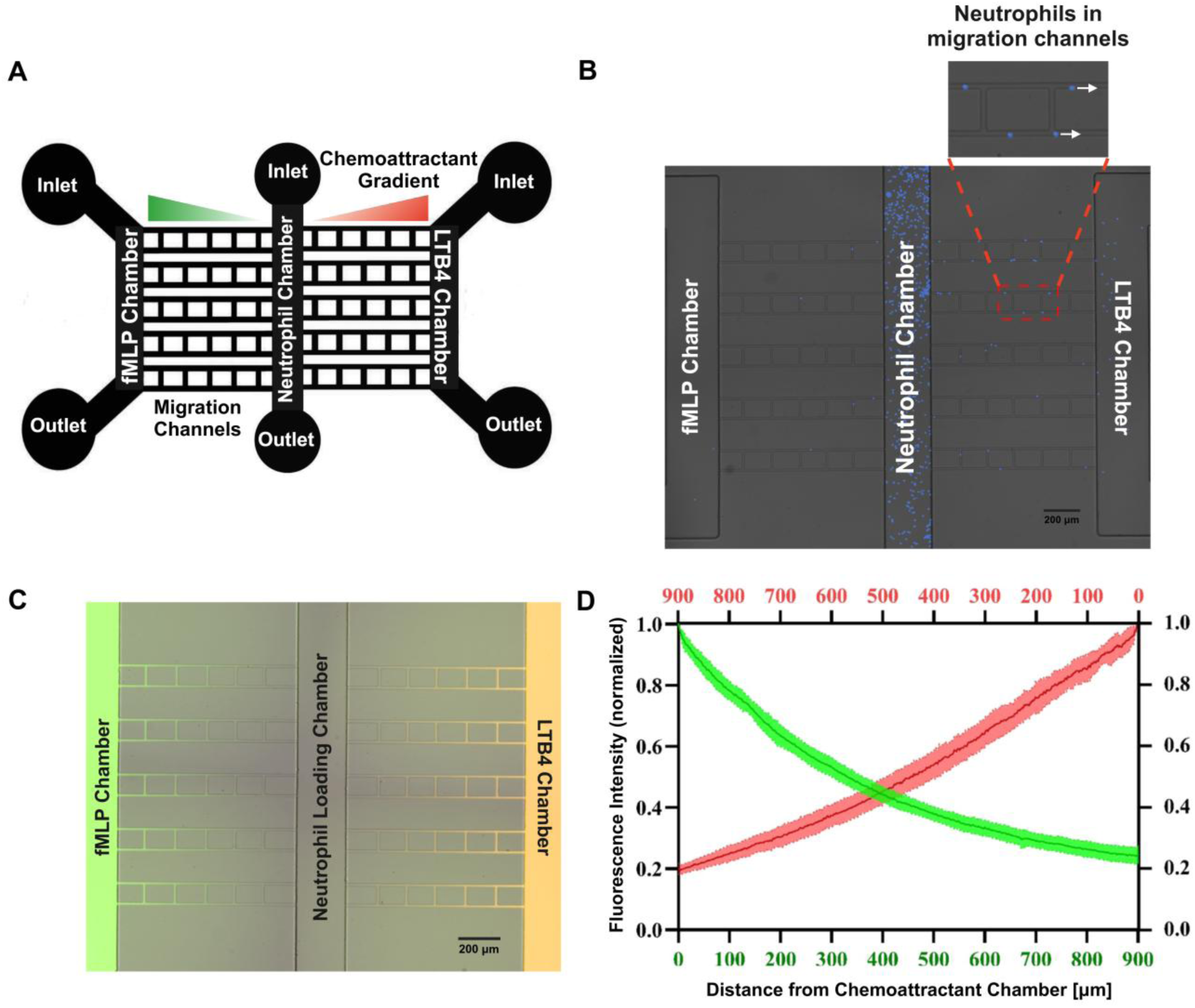
Optimization and validation of microfluidic chemotaxis system to quantify the impact of sex differences and P188 drug treatment on burn-injured murine neutrophil function. (A) Microfluidic device design with two opposing chemoattractant chambers on either side of a central neutrophil chamber with an inlet and outlet port for each of these chambers. The ten perpendicular migration channels connecting the neutrophil chamber to the fMLP and LTB4 chambers on either side are used to study neutrophil chemotaxis to the chemoattractant gradient. (B) Microfluidic chemotaxis system with murine neutrophils in the central chamber, with fMLP in the left chemoattractant chamber and LTB4 in the right chemoattractant chamber. 10X Brightfield and DAPI merged Nikon TiE images. (C) Linear gradient formation was validated in migration channels towards the neutrophil chamber 15 minutes after priming of chemoattractant chambers. FITC-conjugated dextran (MW = 10,000 Da) and TRITC-conjugated dextran (MW = 10,000 Da) image illustrate the linear gradient formation in the left and right migration channels of the microfluidic system, respectively. 10X Brightfield, FITC and TRITC merged images taken using Nikon TiE microscope. (D) Quantification of fMLP and LTB4 gradients formed within the migration channels in the microfluidic chemotaxis system (n= 10 channels). Both fMLP (MW = 438 Da) and LTB4 (MW = 336 Da) have an equal gradient slope.

### 2.7 Microfluidic chemotaxis assay

Three µL [11µg/mL] of fibronectin (Sigma-Aldrich, St. Louis, MO) was injected through the inlets of all three chambers and 40 µL fibronectin was then added on top of all the microfluidic devices to cover all inlets and outlets. The 24 well-plate containing the microfluidic devices was then placed in the vacuum desiccator for 10 minutes, allowing the fibronectin solution to fill all the channels and displace the air from the PDMS devices. After removing the 24-well plate from the desiccator, it was placed at room temperature for at least 30 minutes to allow the fibronectin to absorb into the channels and PDMS surfaces. 1 mL of complete media (IMDM+20% FBS) was then added on top of each well to cover the microfluidic devices inlets and outlets. fMLP (Sigma-Aldrich) and LTB4 (LTB4, Cayman Chemical) chemoattractant solutions were serially diluted in complete media to a final concentration of 100 nM. 4 µL of fMLP [100 nM] and LTB4 [100 nM] chemoattractant solutions were loaded into the left and right chemoattractant channels of the device respectively using gel loading pipette tips. Murine neutrophils at a concentration of 10^7^ cells/mL were then loaded into the central cell loading chamber using a gel loading pipette tip. Finally, the media on top of each device was replaced with new media to remove the residual cells and chemoattractants from each well.

### 2.8 Chemotaxis imaging

Time-lapse imaging experiments were performed at 37◦C with 5% carbon dioxide on a fully automated Nikon TiE microscope, using a Plan Fluor 10x Ph1 DLL (NA = 0.3) lens. Image capture was performed using NIS-elements (Nikon Inc., Melville, NY). Images were recorded using DAPI fluorescent and bright-field channels at four minutes intervals for four hours. Neutrophil migration was quantified as follows: (1) Percentage of neutrophils migrated, (2) velocity of migration, and (3) rate of accumulation.

### 2.9 Quantification of neutrophil migration parameters

Our custom image analysis utilizes the Image Processing Toolbox in MATLAB to detect the neutrophils in the migration channels across all time points and calculate their individual velocities. In specific, our approach applies a Laplacian of Gaussian (LoG) filter to separate the blue color in cells from the background. This is followed by the application of zero-crossings to determine irregular boundaries of the cells. These cells are localized with respect to the migration channels to identify their specific regions. Our approach uses k-nearest neighbors’ algorithm to associate cells between successive time points. A novel implementation in our approach allows for using the area of the cells as the third dimension to keep track of these cells across time points. Additionally, with the knowledge of the time elapsed between two successive images, our approach computes the average velocities of individual neutrophils.

For accumulation rate, straightness and percent neutrophil migration, cells were counted using NIS-Elements software from Nikon. The definitions of migration parameters are listed in Table 1.

**Table 1.** Migration parameter definitions.

| Migratory phenotype | Definition | Units |
| --- | --- | --- |
| Percentage of neutrophils migrated (chemoattractant chamber or migration channels) | Number of Neutrophils in the Chemoattractant chamber or Migration Channels at last Time Point/Total Number of Cells in the Center Loading Channel $\times 100$ | Percentage (%) |
| Rate of accumulation | Slope of 1-hour period that neutrophils accumulated the most in the chemoattractant chamber | % neutrophils accumulated/hour |
| Neutrophil velocity | Distance Traveled by Neutrophils /Time Elapsed | $\mu\text{m}/\text{min}$ |
| Straightness | End Points Distance/Trajectory Length | - |

### 2.10 *Ex vivo* treatment of mouse neutrophils with P188

Isolated neutrophils were treated with 89 µM P188 or complete IMDM control on a 24-well plate for 1 h at 37°C with 5% CO2.

### 2.11 Flow cytometry to examine CD11b expression and ROS production

To assess the activation state of mouse neutrophils in response to the treatments of burn and P188, we performed flow cytometry to measure the expression of neutrophil activation marker CD11b and the production of reactive oxygen species (ROS). After isolation from mouse peripheral blood and ex vivo treatment with P188 where applicable, neutrophils were washed with PBS once and stained with Zombie R685™ Fixable Viability Kit (1:1,000, BioLegend, 423119) to label dead cells for 15 min at room temperature. After washing with Flow Cytometry Staining Buffer (Invitrogen, 00422226) once, cells were treated with TruStain FcX™ PLUS (BioLegend, 156603) for 5 min at 4°C to block nonspecific binding of Fc receptors and stained with the FITC anti-mouse/human CD11b antibody (1:100, BioLegend, 101206) diluted in the staining buffer for 20 min at 4°C. For the ROS production assay, isolated mouse neutrophils were seeded on a V-bottom 96-well plate and treated with 200 nM phorbol 12-myristate 13-acetate (PMA) (Thermo Scientific Chemicals, 356150010) in complete IMDM to induce ROS production for 30 min at 37°C with 5% CO2. Dihydrorhodamine 123 (DHR 123) (Invitrogen, D23806) at 1 µM was pre-added to the mixture to stain ROS. Cells were then incubated for 10 min at 4°C to stop the reaction 1. After washing with the staining buffer once, cell samples were kept on ice in the dark and analyzed by an LSR Fortessa flow cytometer (BD Biosciences) immediately.

Data were analyzed using FlowJo 10.8.1 software (BD Biosciences). For the gating strategy, cells were first gated based on the side scatter area (SSC-A, cell granularity) versus forward scatter area (FSC-A, cell size) density plot to exclude debris. Single cells were then gated based on the forward scatter height (FSC-H) versus forward scatter area (FSC-A) density plot to exclude cell doublets. For CD11b expression, live cells were then gated based on their negative staining for the Zombie dye to exclude dead cells. The median fluorescence intensity (MFI) on the FITC channel was measured to represent the levels of CD11b expression and ROS production in their respective cell samples.

### 2.12 RNA isolation, cDNA synthesis, and ddPCR for FPR1 and BLT1 gene expression analysis

Total RNA was isolated from neutrophils collected from all experimental groups (sham, sham + P188, burn, burn + P188) using the PicoPure™ RNA Isolation Kit (Thermo Fisher Scientific) according to the manufacturer’s instructions. The quality and concentration of the isolated RNA were assessed before proceeding to complementary DNA (cDNA) synthesis. cDNA was synthesized from the RNA samples using the High-Capacity cDNA Reverse Transcription Kit (Applied Biosystems™). The synthesized cDNA was then used as the template for downstream digital droplet PCR (ddPCR) analysis. The ddPCR assay was performed using a QX600 ddPCR system (Bio-Rad), and data were analyzed with QX Manager 2.1 Standard Software. FPR1 and BLT1 expression levels were quantified using gene-specific probe assays with the following configurations: FPR1 (Mice) was detected using the HEX channel, BLT1 (Mice) was detected using the FAM channel, Beta-actin (ACTB) (Mice) served as the reference gene and was detected using the Cy5 channel. Beta-actin was used as a housekeeping gene for normalization to calculate relative expression levels of the target genes. Data were analyzed to compare gene expression levels across all experimental groups.

### 2.13 Statistical analysis

All experiments were conducted and replicated three times. GraphPad Prism software (La Jolla, CA) was used for statistical analysis. Data are presented as mean ± SEM, and significance was determined using ANOVA with Tukey’s post hoc test, and *p* ≤ 0.05 considered statistically significant.

## 3. Results

### 3.1 Optimization and validation of microfluidic chemotaxis system to quantify the impact of sex differences and P188 drug treatment on burn-injured murine neutrophil function

The designed microfluidic chemotaxis system enables the chemoattractant gradient formation from two opposing chemoattractant chambers toward the central cell loading chamber through the long and narrow connecting migration channels (Fig. 1A and B). To mimic the tissue microenvironment, we optimized and evaluated the presence of linear gradient between two opposing chemoattractant chambers toward the central cell loading chamber. We primed the microfluidic chemotaxis device with FITC-conjugated (MW = 10000 Da) dextran in the bacterial chemoattractant (fMLP) chamber and TRITC-conjugated (MW = 10000 Da) dextran in the pro-inflammatory lipid mediator (LTB4) chamber. Here, we added fluorescein (FITC-labeled) dextran to fMLP [100 nM] in the left chamber and added tetramethylrhodamine (TRITC-labeled) dextran to LTB4 [100 nM] in the right chamber, respectively (Fig. 1C). We quantified the formation of gradient in one microfluidic chemotaxis system containing 10 migration channels, and both fMLP (MW = 438 Da) and LTB4 (MW = 336 Da) showed an equal slope for gradient (Fig. 1D).

### 3.2 Visualization of Alexa Fluor–Conjugated P188 in Tissues

Confocal fluorescence microscopy confirmed the presence of Alexa Fluor–conjugated P188 in multiple organs. Fluorescent signal was detected in skin, liver, kidney, lung, and spleen. Nuclear counterstaining with DAPI provided structural context for tissue sections. Representative images are shown in Fig. 2, demonstrating that the conjugated drug was distributed across different organs in both sham and injured animals. These findings qualitatively confirm systemic distribution of P188 following intravenous administration. The images shown were acquired at 60X magnification, while corresponding 20× images are provided in Supplementary Figure 1.

**Figure 2.**
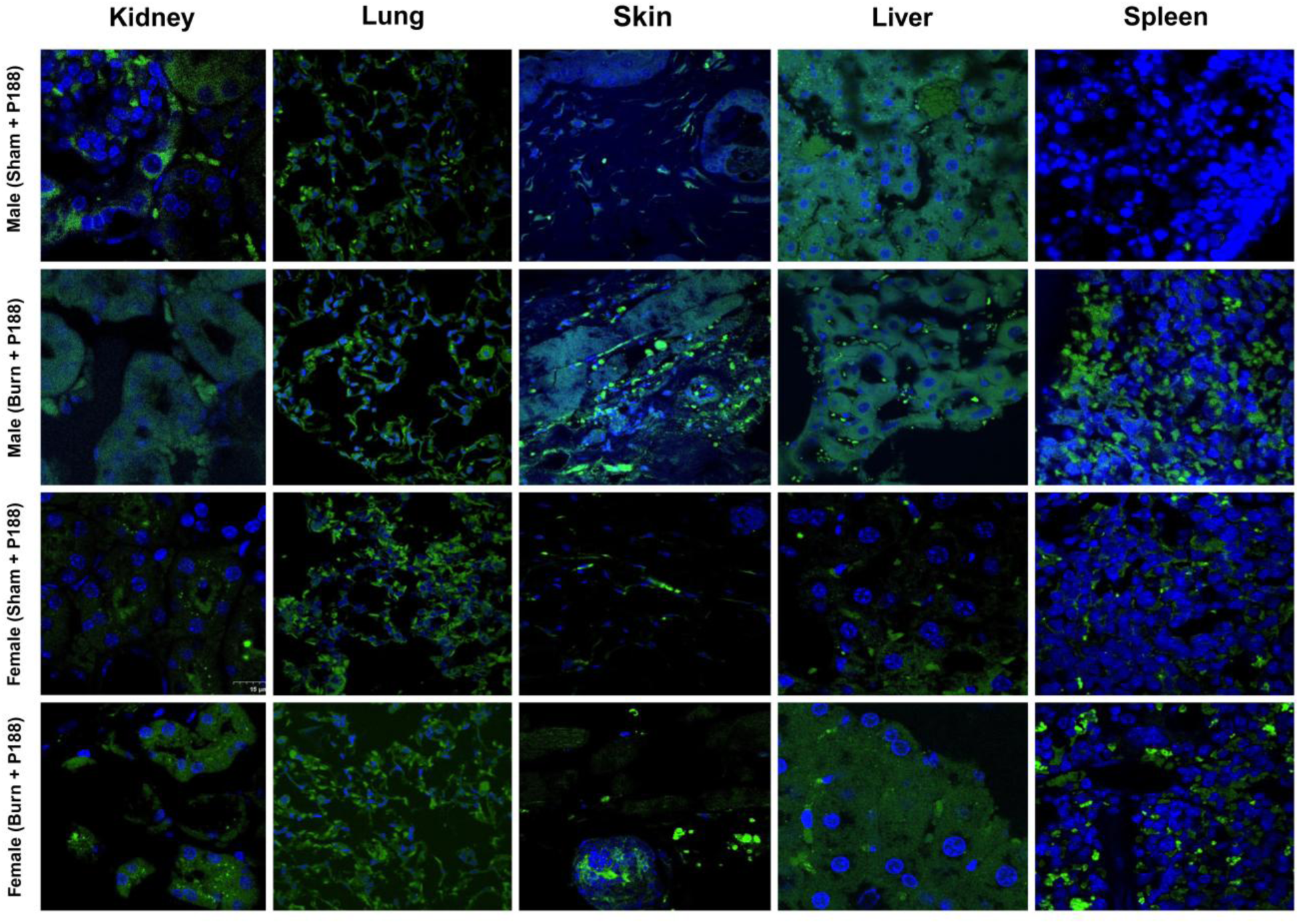
Systemic distribution of Alexa Fluor–conjugated P188 in mouse organs. Representative confocal fluorescence images of skin, liver, kidney, lung, and spleen collected 24 h after intravenous administration of Alexa Fluor–conjugated P188 (15 µg/mouse). Alexa Fluor signal (green) indicates the presence of the conjugated drug, while nuclei are counterstained with DAPI (blue). Images are representative of both sham and burn-injured animals, confirming qualitative systemic distribution of P188. Acquired at 60X magnification.

### 3.3 P188 treatment enhances neutrophil chemotaxis toward LTB4, with a stronger effect observed in females

As shown in Fig. 3 neutrophil migration towards LTB4 was significantly increased in mice given P188 systemically prior to neutrophil isolation (panel A), especially in females (red bars) as opposed to males (blue bars). The statistical significance patterns in panel B differ slightly, suggesting that systemic factors in vivo influence neutrophil responses in ways not fully replicated ex vivo. Burn groups without treatment show decreased neutrophil migration, suggesting that burns impede chemotaxis, while sham groups without P188 display baseline migration levels across both panels. However, P188 treatment significantly improves migration in all groups, with females experiencing a greater impact. According to statistical analysis, P188 increases migration under both sham and burn conditions. For systemic P188 treatment, there are also significant sex-based differences, with females consistently showing a greater migratory response toward LTB4 than males. These results imply that P188 may have therapeutic potential and alleviate burn-induced abnormalities in neutrophil migration. Additionally, sex-based differences in neutrophil behavior highlight the significance of taking biological sex into account when studying immune responses. Table 2 and 3 represent the average migration percentage toward LTB4 [100 nM] among male and female mice. To confirm that the low chemotaxis and migration percentages observed for fMLP are not due to random migration, we conducted control experiments without any chemoattractant. In these negative control conditions, we observed no significant neutrophil migration, as demonstrated in Supplementary Fig 2^12,46^. This indicates that the minimal migration toward fMLP is not attributable to random cell movement.

**Figure 3.**
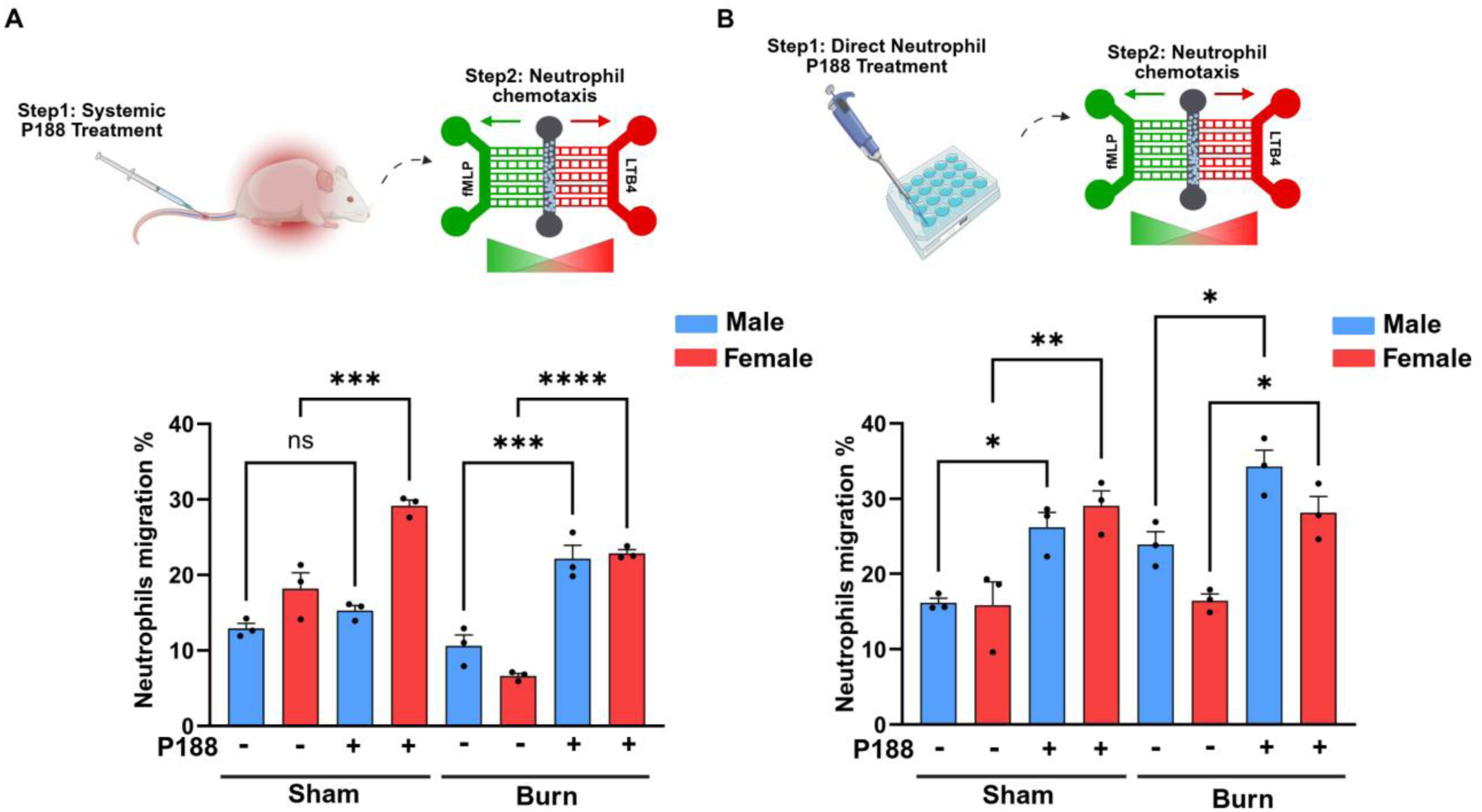
P188 treatment enhances neutrophil chemotaxis toward LTB4, with a stronger effect observed in females. (A) Systemic P188 Treatment, mice were injected with P188, and after 24 hours, neutrophils were isolated from blood and analyzed for migration using a microfluidic system. (B) Direct Neutrophil P188 Treatment, neutrophils were isolated from blood first and then treated with P188 ex vivo for one hour before migration analysis. In both conditions, neutrophil migration percentage toward LTB4 was quantified after four hours, with male (blue bars) and female (red bars) neutrophils analyzed separately. Statistical significance is indicated as follows: *p < 0.05, **p < 0.01, ***p < 0.001, ****p < 0.0001, ns = not significant.

**Table 2.**
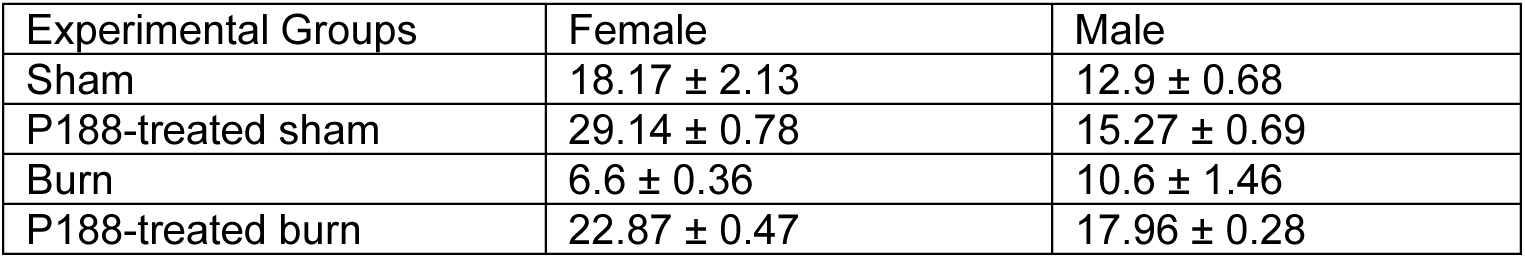
Migration percentage for neutrophils toward LTB4 for Systemic P188 Treatment condition.

**Table 3.** Migration percentage for neutrophils toward LTB4 for Direct Neutrophil P188 Treatment condition.

| Experimental Groups | Female | Male |
| --- | --- | --- |
| Sham | 15.83 ± 3.12 | 16.17 ± 0.62 |
| P188-treated sham | 29.03 ± 2.03 | 26.2 ± 1.97 |
| Burn | 16.47 ± 0.87 | 23.9 ± 1.70 |
| P188-treated burn | 28.13 ± 2.14 | 34.27 ± 2.20 |

**Table 4.** The average neutrophil velocities for the four experimental groups of male and female mice under both systemic P188 treatment and direct neutrophil P188 treatment conditions.

| Experimental group | Systemic P188 Treatment |  | Direct neutrophil P188 treatment |  |
| --- | --- | --- | --- | --- |
| | Male ($\mu\text{m}/\text{min}$) | Female ($\mu\text{m}/\text{min}$) | Male ($\mu\text{m}/\text{min}$) | Female ($\mu\text{m}/\text{min}$) |
| Sham | 6.827 ± 0.830 | 6.050 ± 0.510 | 6.071 ± 0.344 | 4.926 ± 0.105 |
| Sham treated with P188 | 7.491 ± 0.454 | 6.910 ± 0.322 | 5.006 ± 0.412 | 4.678 ± 0.604 |
| Burn | 9.897 ± 0.706 | 9.561 ± 0.319 | 6.863 ± 0.339 | 6.484 ± 0.691 |
| Burn treated with P188 | 7.571 ± 0.412 | 6.732 ± 0.382 | 5.850 ± 0.634 | 4.490 ± 0.327 |

### 3.4 P188 treatment mitigates burn injury-induced increases in neutrophil velocity, with sex-dependent differences

For better understanding of neutrophil migratory behaviors, we measured single-cell velocities of neutrophils from four different groups of male and female mice migrating toward LTB4 [100 nM] chemoattractant gradient (Fig. 4). The average velocities of female murine neutrophils towards LTB4 in the sham, P188-treated sham, burn, and P188-treated burn groups for systemic P188 treatment are 6.050 μm/min, 6.910 μm/min, 9.561 μm/min, and 6.732 μm/min, respectively (Fig. 4A). The velocities of male murine neutrophils towards LTB4 for systemic P188 treatment condition in the experimental groups, listed in the same order, are 6.827 μm/min, 7.491 μm/min, 9.897 μm/min, and 7.571 μm/min, respectively (Fig. 4A). The average velocities of female murine neutrophils towards LTB4 in the sham, P188-treated sham, burn, and P188-treated burn groups for direct neutrophil P188 treatment condition are 4.926 μm/min, 4.678 μm/min, 6.484 μm/min, and 4.490 μm/min, respectively (Fig. 4B). The velocities of male murine neutrophils towards LTB4 for direct neutrophil P188 treatment condition in the experimental groups, listed in the same order, are 6.071 μm/min, 5.006 μm/min, 6.863 μm/min, and 5.850 μm/min, respectively (Fig. 4B).

**Figure 4.**
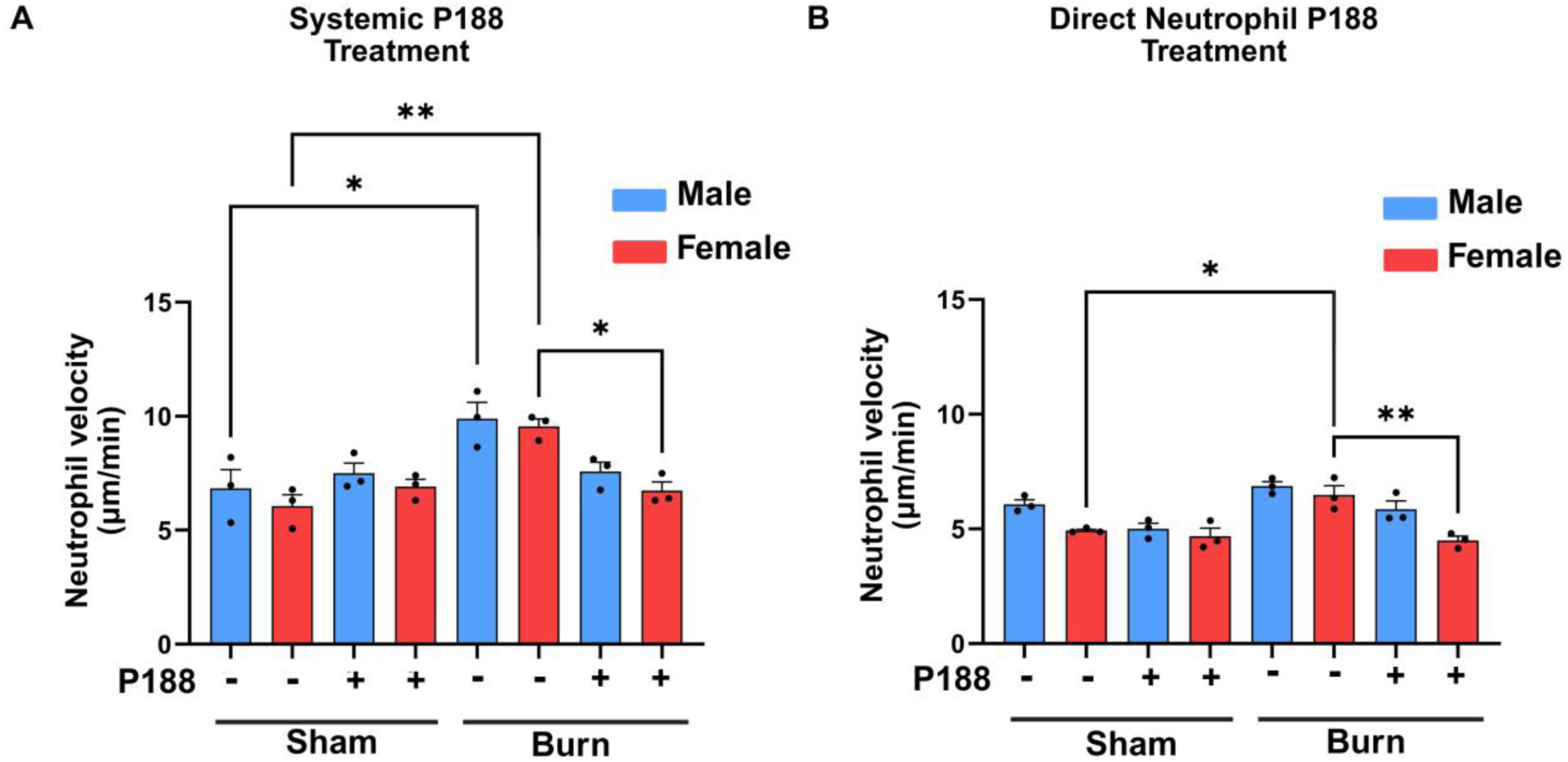
P188 treatment mitigates burn injury-induced increases in neutrophil velocity, with sex-dependent differences. In panel A, systemic P188 treatment was administered in vivo before blood collection, while in panel B, neutrophils were isolated first and then treated with P188 ex vivo. Systemic P188 treatment alleviated the burn injury-induced increase in neutrophil velocity, bringing it closer to sham levels, while direct P188 treatment increased neutrophil velocity in both sham and burn groups. Statistical significance is indicated by *p < 0.05 and **p < 0.01.

In both male and female mice, neutrophils from burn conditions exhibit the highest migration velocity, indicating a significant increase in motility following burn injury compared to sham controls. Interestingly, P188 treatment in burn mice partially reduces neutrophil velocity. The average neutrophil velocities towards LTB4 chemoattractant for the four experimental groups of male and female mice are shown in table 5.

**Table 5.** Neutrophils accumulation rate for Systemic P188 Treatment.

| Experimental group | LTB4 (%) |  |
| --- | --- | --- |
|  | Male | Female |
| Sham | 21.333 ± 1.841 | 14.767 ± 0.233 |
| Sham treated with P188 | 29.433 ± 2.979 | 30.033 ± 5.053 |
| Burn | 24.500 ± 1.762 | 22.167 ± 1.819 |
| Burn treated with P188 | 27.33 ± 1.338 | 35.367 ± 1.994 |

### 3.5 P188 increased the accumulation rate for sham and burn-injured female neutrophils

As shown in Fig. 5A (Systemic P188 Treatment), P188 treatment significantly increased neutrophil accumulation rates in female mice, indicating that systemic P188 enhances neutrophil adhesion or migration in females following both sham and burn conditions. In contrast, male neutrophils showed a limited response to systemic P188 treatment, with no significant increase in accumulation rates. This suggests a potential sex-specific difference in how neutrophils respond to systemic P188 treatment, where females exhibit a stronger pro-adhesive effect. In panel B (Direct Neutrophil P188 Treatment), direct P188 treatment significantly increased neutrophil accumulation rates in sham conditions for both males and females. However, in burn-injured groups, P188 treatment had no significant effect on accumulation rates, suggesting that burn-induced changes in neutrophil behavior may counteract any enhancing effects of P188 treatment. Overall, these findings suggest that systemic P188 treatment selectively enhances neutrophil accumulation in females, whereas direct P188 treatment increases accumulation only in sham conditions for both sexes. The neutrophil accumulation rates for all conditions are shown in Tables 5 and 6.

**Figure 5.**
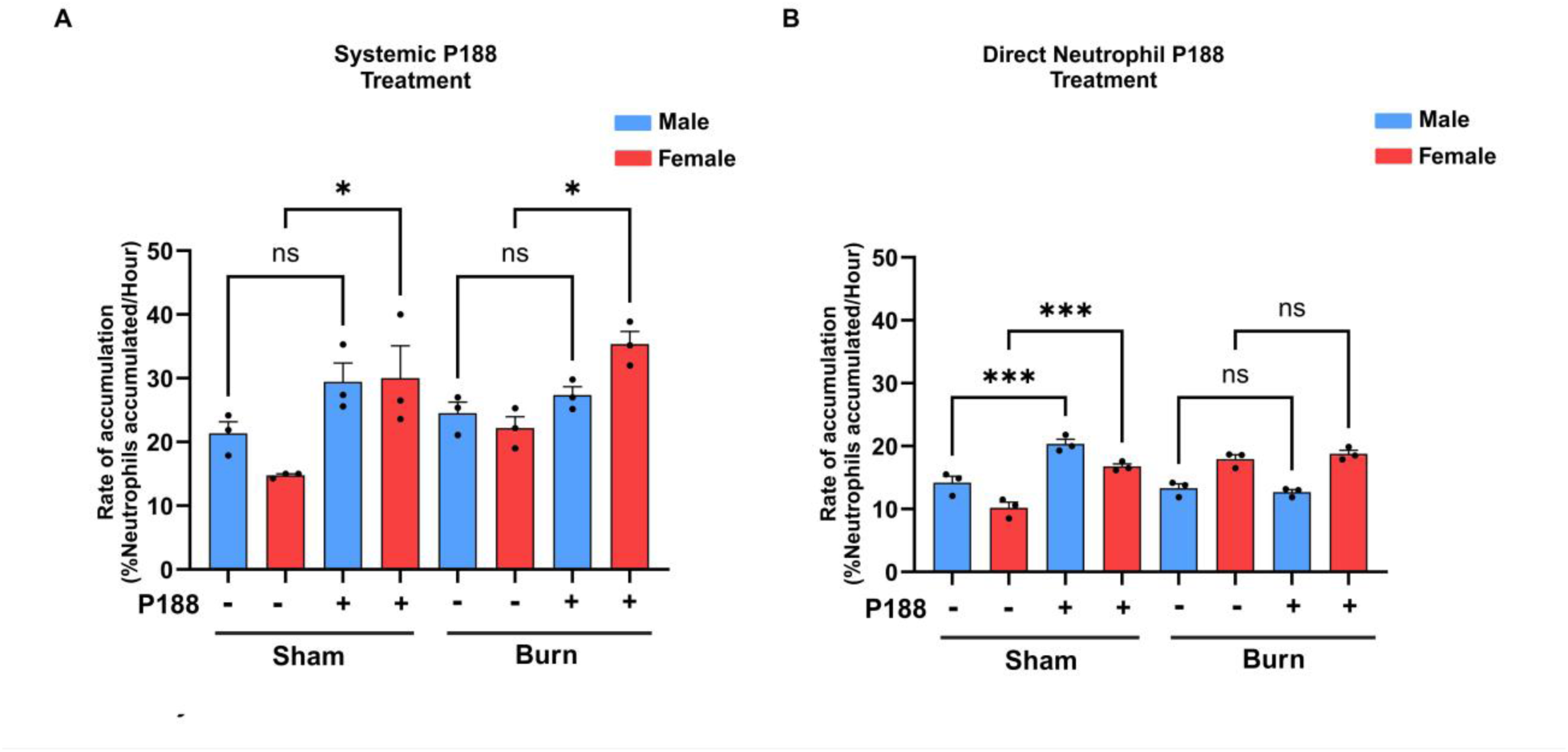
P188 increased the accumulation rate for sham and burn-injured female neutrophils. (A) Systemic administration of P188 increased neutrophil accumulation in both male and female sham groups compared to untreated controls (p < 0.05), but it had no significant effect on neutrophil accumulation in burn groups. (B) Direct Neutrophil P188 Treatment: Unlike systemic treatment, direct ex vivo P188 treatment significantly increased neutrophil accumulation in sham groups for both males and females (p < 0.001), while having no impact on burn groups, similar to systemic treatment. Statistical significance: ns (not significant), *p < 0.05, ***p < 0.001.

**Table 6.** Neutrophils accumulation rate for Direct Neutrophil P188 Treatment.

| Experimental group | LTB4 (%) |  |
| --- | --- | --- |
|  | Male | Female |
| Sham | 14.167 ± 1.048 | 10.200 ± 0.889 |
| Sham treated with P188 | 20.367 ± 0.745 | 16.167 ± 0.203 |
| Burn | 13.300 ± 0.709 | 17.933 ± 0.717 |
| Burn treated with P188 | 12.700 ± 0.404 | 18.767 ± 0.593 |

### 3.6 Systemic P188 treatment enhances neutrophil directional migration post-burn

In Fig. 6 panel A, systemic P188 treatment significantly decreases the directionality of neutrophils in burn-injured mice for both males and females, indicating that P188 negatively impacts neutrophil migration in this condition. Additionally, female neutrophils exhibit generally lower directional migration compared to male neutrophils under systemic P188 treatment, suggesting a sex-dependent difference in migratory behavior. In contrast, panel B shows that direct P188 treatment does not significantly alter neutrophil straightness in burn-injured mice, highlighting the importance of in vivo systemic mechanisms in mediating the effects of P188 on neutrophil migration. The violin plots in panels C and D represent the single cell neutrophil distribution for directional migration. These findings emphasize that systemic administration of P188 plays a crucial role in altering neutrophil migration, while direct exposure alone is not sufficient to induce significant changes.

**Figure 6.**
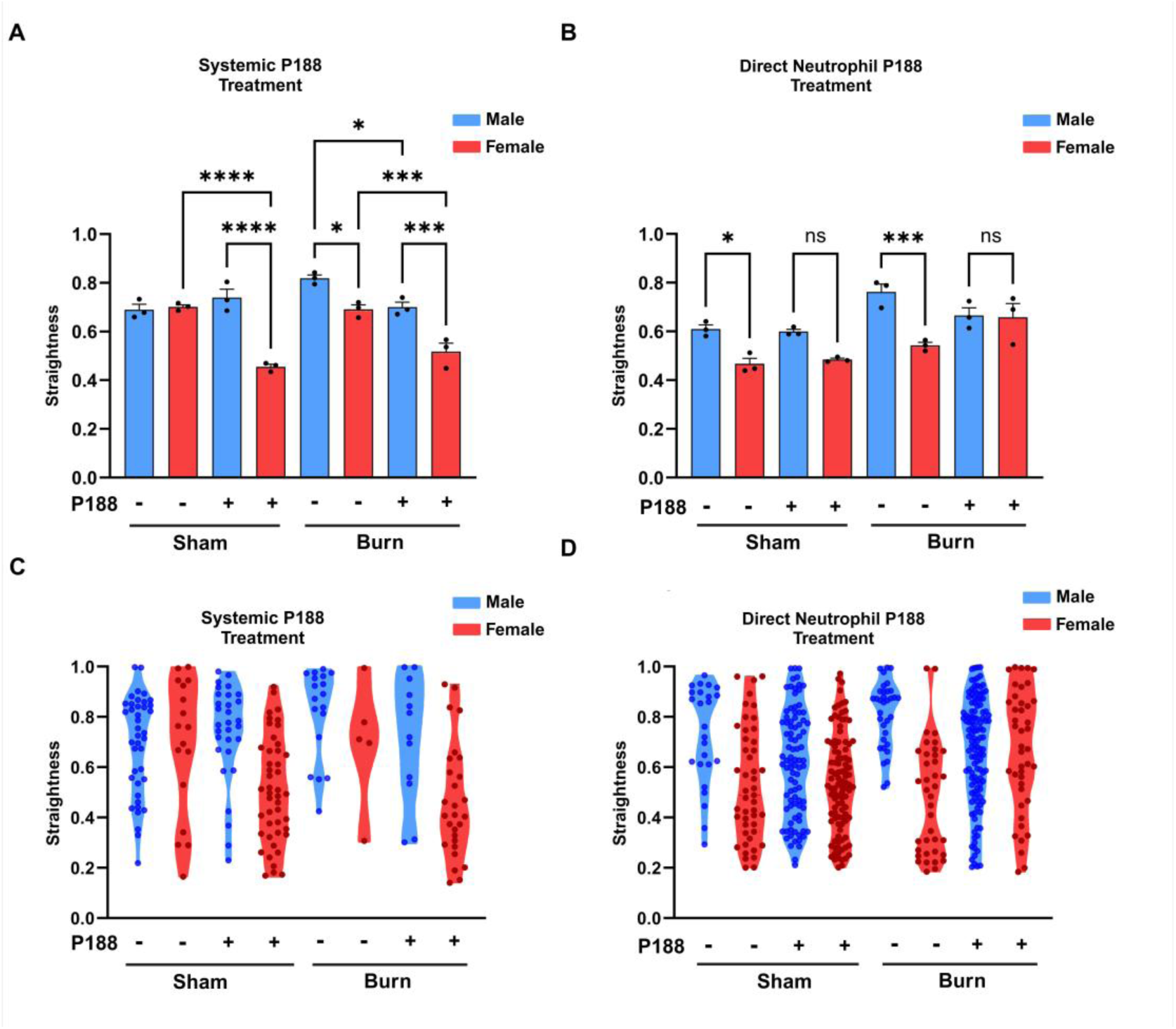
Systemic P188 treatment enhances neutrophil directional migration post-burn. Figure investigates the effects of burn injury and P188 treatment on neutrophil migratory straightness in male and female mice under both systemic P188 treatment (A, C) and direct neutrophil P188 treatment (B, D). Systemic P188 treatment significantly decreases the directionality of neutrophils in burn-injured mice for both males and females, indicating that P188 negatively impacts neutrophil migration in this condition.

### 3.7 Burn injury induces upregulation of FPR1 and BLT1 in neutrophils

The data presented in Fig. 7A show the relative expression of FPR1 in neutrophils from male and female mice across different treatment groups. In the sham condition, systemic P188 treatment does not significantly alter FPR1 expression in either sex. However, in the burn-injured group, there is a significant increase in FPR1 expression compared to sham, with P188 treatment not significantly affecting this elevated expression level. Interestingly, while both male and female burn-injured mice show an upregulation of FPR1, no significant sex-based differences are observed. In Fig. 7B the BLT1 expression shows a different trend. BLT1 expression shows no difference between sham and P188-treated sham groups. However, burn injury significantly increases BLT1 expression compared to sham, and P188 treatment does not significantly alter this elevated expression level in burn-injured mice. These findings indicate that burn injury induces upregulation of both FPR1 and BLT1 in neutrophils, which may contribute to enhanced chemotactic signaling in the inflammatory response. This highlights the complexity of P188’s role in modulating neutrophil function, suggesting that inflammatory conditions such as burn injury may diminish any potential regulatory effects of P188 on these chemotactic receptors. Further studies are necessary to study the molecular mechanisms behind these observations, particularly in the context of systemic vs. localized P188 effects and potential sex-specific immune responses. No significant changes were observed in FPR1 and BLT1 expressions following direct neutrophil P188 treatment (Supplementary Fig. 3). We measured CD11b expression and ROS production in mouse neutrophils, and the data from these experiments are provided in Supplementary Figure 4.

**Figure 7.**
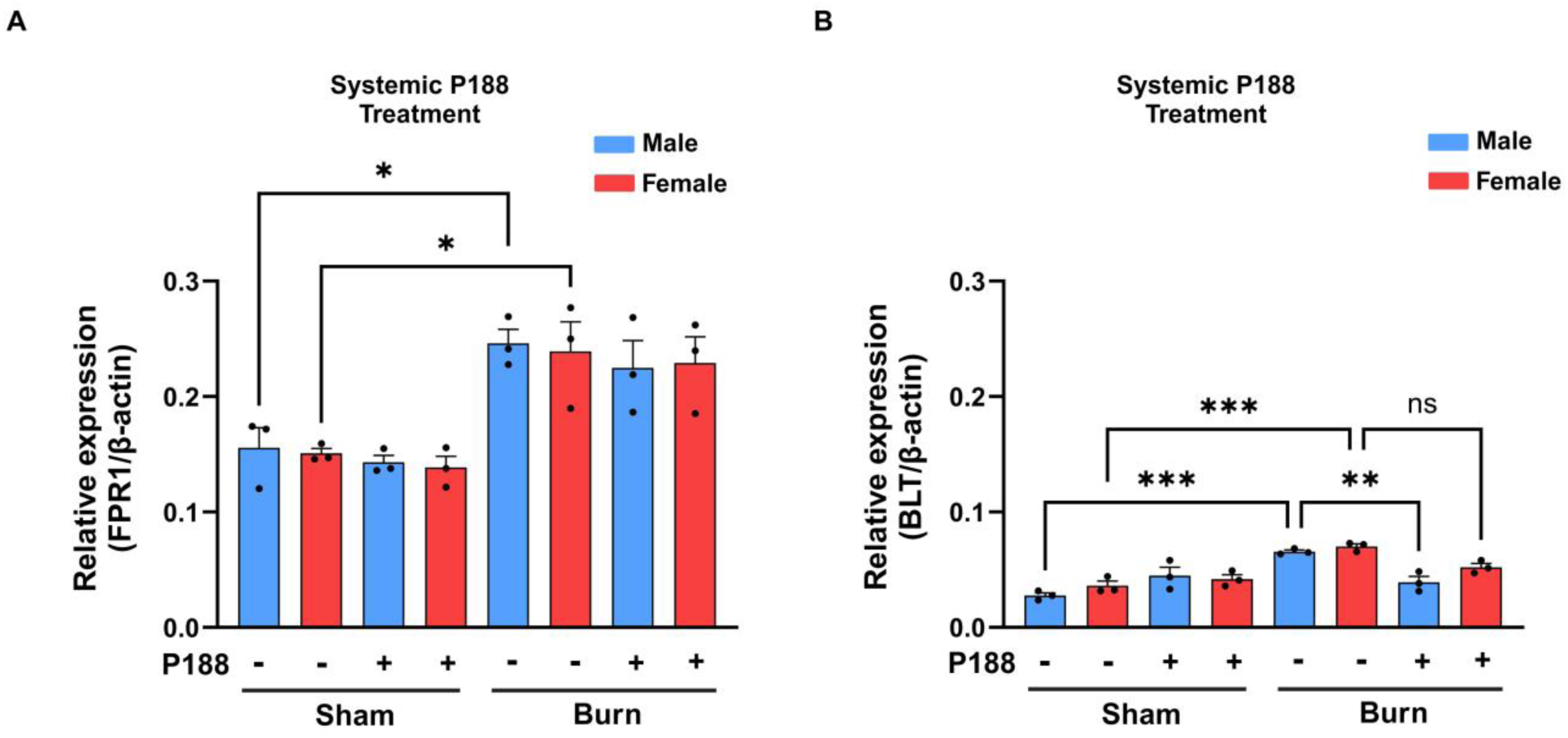
Burn injury induces upregulation of both FPR1 and BLT1 in neutrophils expression level. Expression levels of FPR1 (A) and BLT1 (B) in neutrophils from male (blue) and female (red) mice under different conditions: sham, sham + P188, burn, and burn + P188. Data are normalized to β-actin expression. Systemic P188 treatment was administered in both sham and burn conditions. Statistical significance is indicated as follows: *p < 0.05, **p < 0.01, ***p < 0.001, and ns (not significant). Error bars represent standard error of the mean (SEM).

## Discussion

Our findings uncover a previously unrecognized role of P188 in modulating neutrophil chemotaxis after burn injury. Our study demonstrates differences in neutrophil chemotaxis following burn injury and treatment P188 using a microfluidic system in systemic P188 treatment (in vivo) and direct neutrophil P188 treatment (ex vivo) conditions. The results show that P188 effectively modulates neutrophil migration, enhancing neutrophil migration toward LTB4 in both male and female mice for systemic P188 treatment (in vivo) and direct neutrophil P188 treatment (ex vivo) conditions **(Fig. 8)**. This indicates a shift in neutrophil preference towards the pro-inflammatory lipid mediator LTB4 over the bacterial chemoattractant, fMLP, suggesting that P188 may promote a more resolving inflammatory response. We observed a significant increase in the velocity of neutrophils from burn mice, in both male and female groups, for systemic P188 treatment (in vivo) and direct neutrophil P188 treatment (ex vivo) conditions, which restored after P188 treatment. This highlights the P188 drug potential to improve dysfunctional migration of neutrophils, resulting in a better and faster immune response and enhanced pathogen clearance, especially in females.

**Figure 8.**
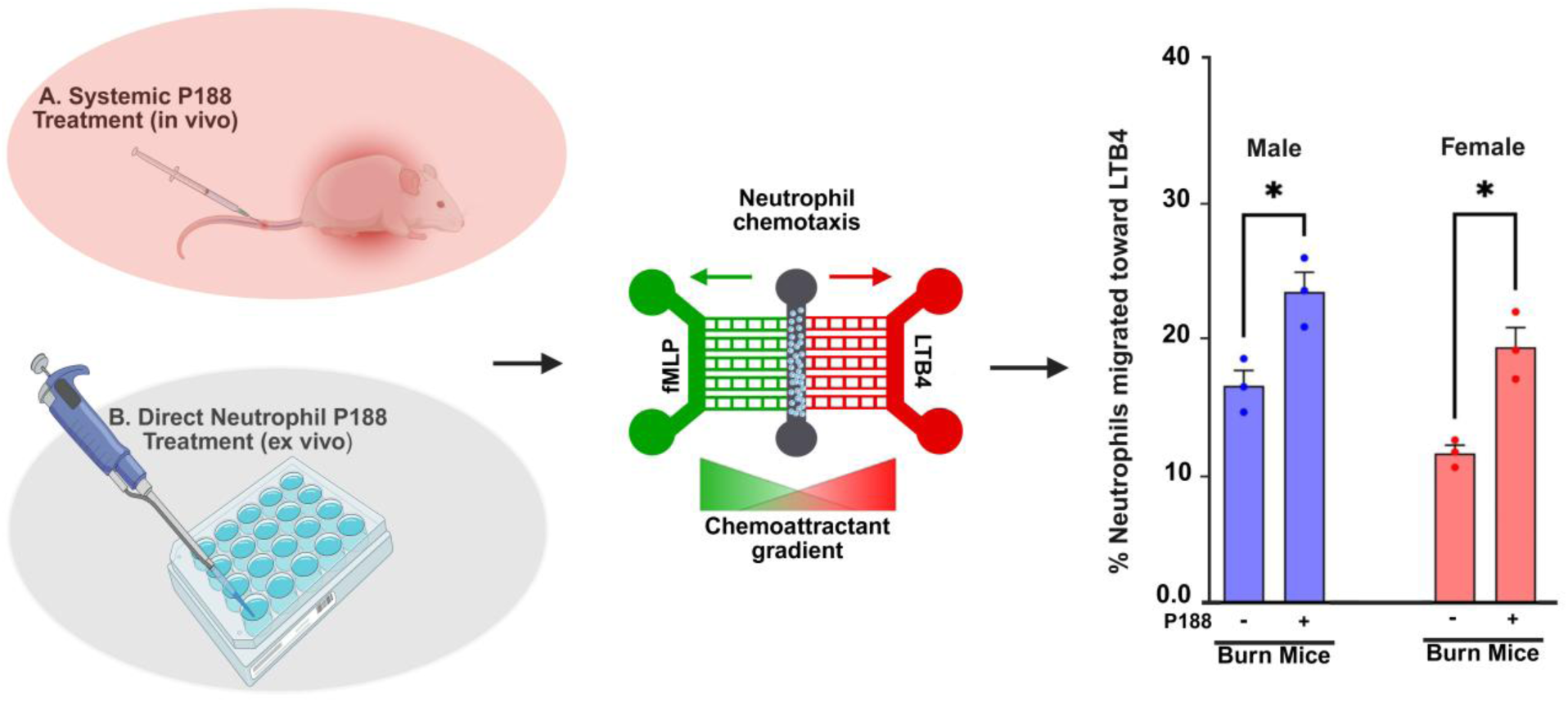
Effects of systemic and direct neutrophil P188 treatment on chemotactic migration in burn-injured mice. This schematic illustrates the experimental approach for evaluating neutrophil chemotaxis in burn-injured mice following two different P188 treatment methods. (A) Systemic P188 treatment (in vivo) involves injecting P188 into burn-injured mice. (B) Direct neutrophil P188 treatment (ex vivo) applies P188 directly to isolated neutrophils in vitro. Neutrophil migration was assessed using a microfluidic system with an fMLP-LTB4 chemoattractant gradient. The bar graph on the right quantifies neutrophil migration toward LTB4 in male (blue) and female (red) burn-injured mice, showing a significant increase in migration following P188 treatment (p < 0.05).

FPR1 (Formyl Peptide Receptor 1) and BLT1 (Leukotriene B4 Receptor 1) are key regulators of neutrophils’ migration and activation during inflammation. Our finding shows the importance of FPR1 and BLT1 expressions in burn injury and P188 treatment. Based on the results, we found that for systemic P188 treatment and FPR1 expression levels, there was a significant increase when burn injuries were introduced to mice. This increase was consistent across both male and female conditions (Fig. 7). In Figure. 7B the BLT1 expression shows a different trend. BLT1 expression shows no difference between sham and P188-treated sham groups. However, burn injury significantly increases BLT1 expression compared to sham, and P188 treatment does not significantly alter this elevated expression level in burn-injured mice. These findings indicate that burn injury induces upregulation of both FPR1 and BLT1 in neutrophils, which may contribute to enhanced chemotactic signaling in the inflammatory response. For FPR1, no significant differences were observed between male and female groups or between systemic (in vivo) and direct neutrophil (ex vivo) P188 treatment conditions.

This suggests that there is an in vivo factor affecting and influencing the marker’s expression and regulatory functions. The fact that P188 did not change the expression of either BLT1 or FPR1 in both systemic P188 treatment (in vivo) and direct neutrophil P188 treatment (ex vivo) conditions implies that its mechanism of action may not directly involve these pathways. We hypothesized that P188 may apply its modulatory effects through other molecular mechanisms which are not directly dependent on the expression of these specific chemotactic receptors. Based on the study findings, targeting BLT1-mediated pathways could be an important strategy in improving immune response and neutrophil chemotaxis following burn injury. However, P188’s therapeutic effects might involve alternate pathways beyond FPR1 and BLT1 regulation.

In general, we described the effects of sex and the drug P188 on the migratory behavior of immune cells and their activation by measuring FPR1 and BLT1 which are chemotactic receptors. When neutrophils from P188-treated burn-injured mice were exposed to the bacterial chemoattractant, fMLP and pro-inflammatory lipid mediator, LTB4 chemoattractant, they showed different migratory behaviors. The neutrophil migration toward LTB4 significantly increased compared to neutrophils from untreated burn mice, in both male and female groups among systemic P188 treatment and direct neutrophil P188 treatment conditions (Fig. 3).

Further research is needed to explore the potential clinical implications of P188 treatment for enhancing immune function and recovery following burn injuries. Additionally, expanding the study to include other immune cell types and their interactions within the inflammatory milieu will help to create a more comprehensive understanding of the immune response to burn injuries. These studies should include mouse models and, eventually, clinical studies to validate the efficacy and safety of P188 in human patients. In this context, it is important to examine how P188 has been applied in existing clinical studies. Regarding the clinical study of P188, it has primarily been used topically in burn injury models and not delivered systemically^47^. The only other clinical studies involving P188 are in sickle cell crisis and during stenting procedures for coronary artery disease, all with the premise that P188 was originally approved for the use as packed red blood cell storage to decrease the viscosity^48,49^. Hence, the only clinical applications are to promote blood flow, not minimize membrane damage, as the other preclinical studies have started to show^16,47^.

Moreover, exploring combination therapies that include P188, and other anti-inflammatory agents may enhance the overall therapeutic effect and provide better outcomes for burn patients. Finally, advancements in microfluidic technology will be leveraged to develop more sophisticated models that can mimic the complex tissue environment and multicellular interactions seen in vivo. Integrating models that include 3D human tissue microenvironments and vascularization with high-throughput screening techniques will accelerate the identification of new therapeutic targets and the development of personalized treatment strategies for burn and other inflammatory conditions. By continuing to investigate the chemotactic effects of treatments such as P188, we move toward more effective and personalized strategies for burn injury management and improved patient care.

## Acknowledgments and sources of funding

The in vivo part of this study was supported by NIH R35 GM138020, and the chemotaxis part of the research was supported by NIH NIGMS (5R35GM133610-02).

## Authorship contribution statement

HRB, and CNJ conceived and designed all the experiments. HRB and ES performed all the microfluidic chemotaxis experiments. HRB, USD, and BV performed image processing and data analysis. HRB and SS conducted the ex vivo P188 treatment, followed by flow cytometry analysis and ddPCR assays. MI, RK, NA, and HRB contributed to the fluorescent imaging experiments involving P188. HRB, CNJ and VN supervised general work. All authors contributed to the article and approved the submitted version. All authors assisted in writing and editing the manuscript. NA, RK, KM, and RW performed the systemic P188 drug administration and handled blood drawings from different groups of mice.

## Supplementary material

Supplementary materials are available at Journal of Leukocyte Biology online.

